# The Nestin neural enhancer is essential for normal levels of endogenous Nestin in neuroprogenitors but is not required for embryo development

**DOI:** 10.1101/2021.03.25.436942

**Authors:** Ella Thomson, Ruby Dawson, Chee Ho H’ng, Fatwa Adikusuma, Sandra Piltz, Paul Q Thomas

## Abstract

Enhancers are vitally important during embryonic development to control the spatial and temporal expression of genes. Recently, large scale genome projects have identified a vast number of putative developmental regulatory elements. However, the proportion of these that have been functionally assessed is relatively low. While enhancers have traditionally been studied using reporter assays, this approach does not characterise their contribution to endogenous gene expression. We have studied the murine Nestin (*Nes)* intron 2 enhancer, which is widely used to direct exogenous gene expression within neural progenitor cells in cultured cells and *in vivo*. We generated CRISPR deletions of the enhancer region in mice and assessed their impact on *Nes* expression during embryonic development. Loss of the *Nes* neural enhancer significantly reduced *Nes* expression in the developing CNS by as much as 82%. By assessing NES protein localization, we also show that this enhancer region contains repressor element(s) that inhibit *Nes* expression within the vasculature. Previous reports have stated that *Nes* is an essential gene, and its loss causes embryonic lethality. We also generated 2 independent *Nes* null lines and show that both develop without any obvious phenotypic effects. Finally, through crossing of null and enhancer deletion mice we provide evidence of *trans*-chromosomal interaction of the *Nes* enhancer and promoter.

## Introduction

Embryonic development requires precise coordinated expression of thousands of genes across space and time. Regulatory elements such as enhancers have a critical role in coordinating spatio-temporal gene expression during embryogenesis. Enhancers are typically located within introns and intergenic regions and comprise DNA motifs that can be bound by transcription factors (TF). TF binding promotes interaction of the enhancer with the target promoter via DNA looping. This process, which involves cohesins and the mediator complex (1), allows TF-associated co-activators to engage the transcriptional machinery and stimulate RNA Pol II-mediated transcription of the target gene. While enhancers are generally regarded to function as *cis*-acting elements, recent evidence suggests that some enhancers can act in *trans* to influence expression of their target gene on the homologous chromosome. *Trans* enhancer-promoter interaction in *Drosophila*, termed transvection, is relatively well characterised and has recently been visualised within developing embryos (2). Few examples of *trans* interactions have been reported in vertebrates, although a recent analysis at the IGH super-enhancer indicates that *trans*-enhancer activity can occur in mammals (3).

The Nestin gene (*Nes*) encodes an intermediate filament protein and is widely expressed during embryonic development including progenitor cells throughout the neuroaxis (4, 5). Differing reports of NES functionality have been published, with Mohseni, Sung (6) suggesting *Nes* is not essential for development of the central nervous system, in contrast to an earlier paper (7) indicating that loss of the gene results in embryonic lethality. The *Nes* neural enhancer (8) is a highly conserved element located in intron 2 and is commonly used to drive exogenous gene expression in neural progenitor cells *in vivo* and *in vitro* (9-11). *In vitro* and transgenic data indicate that TFs belonging to the SOX and POU families bind the *Nes* enhancer and function synergistically to control the *Nes* expression in the CNS progenitors (12). Consistent with these data, ChIP-seq experiments have identified robust binding of endogenous SOX3 protein at the *Nes* enhancer in cultured neuroprogenitor cells (13).

Traditionally, enhancers have been identified and characterized using transgenic reporter assays (14). This has proven to be a useful approach to determine the contribution of specific enhancer elements to the spatiotemporal expression of its cognate gene. However, this strategy is incapable of recapitulating the endogenous genomic and chromatin environment in which the enhancer is usually located. CRISPR gene editing technology (15) enables rapid and efficient deletion of target sequences *in vivo*, providing a valuable tool to assess putative enhancer function in an endogenous context. This is an important advancement as the number of putative enhancers identified via bioinformatic and TF binding studies continues to grow, while functional studies are lagging.

Despite widespread use of the *Nes* neural enhancer, the contribution of this enhancer to *Nes* expression during development has not been studied, nor have the effects of removing the enhancer on the developing CNS. Here we show that CRISPR-mediated deletion of the *Nes* enhancer results in a significant reduction in mRNA expression as well as altered protein levels within the developing mouse central nervous system. Using CRISPR/Cas9, we also generate two *Nes* loss of function mouse lines and show that *Nes* KO mice are viable. Finally, we present evidence that the *Nes* enhancer is able to function in *trans*.

## Materials and Methods

### Mouse Generation

Mutant mice were generated by CRISPR microinjection as previously described(16, 17). In brief, CRISPR guides were designed using the crispr.mit.edu tool to determine off-target scores. Guide RNA sequences (*Nes* enhancer deletion-TTTGCGGTCTGAAAAGGATT, AGAATCGGCCTCCCTCTCCG, *Nes* null lines - GGAGCTCAATCGACGCCTGG, GCACAGGAGACCCTACTAAA) were annealed and ligated into px330 (Addgene, #42230) after digestion with *BbsI* (NEB) using Rapid Ligation Kit (ThermoFisher Scientific), and transformed into *E. Coli* using standard protocols. Plasmid was extracted from positive colonies using a Midi-Prep kit (Qiagen). Primers were designed to incorporate T7 promoter sequence and tracR sequence, and PCR was performed on plasmid DNA with Phusion High Fidelity PCR Kit (NEB). PCR products were converted to RNA using the T7 RNA Transcription Kit (NEB) and purified with RNEasy Kit (Qiagen) to generate sgRNA. Cas9 mRNA was synthesised from the *XhoI* (NEB) digested pCMV/T7-hCas9 (Toolgen) using the Mmessage Mmachine T7 Ultra Transcription Kit (ThermoFisher).

BL6/2J females were superovulated with Pregnant Mare Serum Gonadotropin (PMSG) and human Chorionic Gonadotropin (hCG) prior to mating with BL6 males for zygote harvesting. Single cell zygotes were collected on the day of microinjection and treated with hyaluronidase to remove surrounding cumulus cells. Cytoplasmic injection was performed with CRISPR reagents (100ng/µL Cas9 mRNA, 50ng/µL sgRNA) before transfer into psuedopregnant CD1 females.

Genomic DNA was extracted from 3 week old tail or ear biopsies using KAPA Mouse DNA Extraction Kit (KAPA Biosystems) or High Pure PCR Template Kit (Roche).

Founder mice were genotyped using FailSafe PCR Kit (EpiCentre) and run on a 12% polyacrylamide gel for heteroduplex assay. The genotype of the founder mice was confirmed via Sanger sequencing after BigDye Terminator v3.1 (Applied Biosystems) PCR reaction using reverse primer.

Regular colony and embryo genotyping was performed with primers flanking deleted sequence (enhancer deletion line F-GCCCCAGTCAGTCTTCTGAG R-GCCACTGCAGGATCACTCTT, *Nes* null FS F1– CTGCTGAGCTGGGATGATGC F2– AGCTCAATCGACGCCTGGA R- GCATTCTTCTCCGCCTCGA, *Nes* null BD F-CTGCTGAGCTGGGATGATGC R- CTGCTGAGCTGGGATGATGC) using 2G Fast MasterMix (KAPA), or Buffer J (EpiCentre) with Taq Polymerase (Roche).

All mouse breeding and experimental work was performed at the University of Adelaide in accordance with relevant ethics approvals (S-201-2013 and S-173-2015).

### Tissue Preparation

Heterozygous (WT/-255) males and females were time mated for embryo collection. Females were humanely killed via cervical dislocation and embryos removed and stored in cold 1x PBS until dissected. Tails were removed and kept at -20ºC. Heads were removed and flash frozen on dry ice and kept at -80ºC for RNA extraction or kept overnight in 4% paraformaldehyde in PBS, washed 3x in PBS and cryoprotected overnight in 30% sucrose before flash freezing in OCT and stored at -20ºC for immunohistochemical analysis.

### Immunohistochemistry

Trunks were sectioned at 16µm on a cryostat (Leica CM1900) and slides washed 3x 10mins in PBT (1xPBS, 0.25% Triton-X), blocked for 30min in Blocking Solution (1x PBS, 0.25% Triton-X, 10% Horse Serum) and then stained overnight with 20µL primary antibody diluted in Blocking Solution and kept in a humidified chamber at 4ºC. Slides were washed 3 x for 10mins in PBS. 200 µL of secondary antibody diluted in Blocking Solution was added to the slides and incubated in a dark humidified chamber for 4hrs at RT. Slides were washed 3 x for 10mins in PBS, dried, mounted with Prolong Gold Antifade + DAPI (Molecular Probes) and coverslipped. Slides kept overnight in the dark before image acquisition using a Nikon Eclipse Ti Microscope using ND2 Elements software. Images were modified for colour, brightness and contrast using Adobe Photoshop v7 (Adobe Systems). Antibodies used were Anti-SOX3 (R&D Systems, AF2569, 1/200), Anti-Nestin (Abcam AB82375, 1/1000), Anti-CD31 (BD Pharmingen 550274, 1/100). Secondary antibodies, Donkey anti-Goat-Cy3 (Jackson ImmunoResearch, 1/400), Donkey anti-Rat-Cy5 (Jackson ImmunoResearch, 1/400), Donkey anti-Rabbit-488 (Jackson ImmunoResearch, 1/400).

### *In situ* Hybridization

*In situ* hybridization probes were designed to target exon 4 of the *Nes* gene. Primers corresponding to the region of interest were used to PCR amplify WT mouse cDNA and incorporate a T7 promoter at the 5’ end. The PCR product was transcribed using the T7 IVT Kit (NEB), followed by DNase I (NEB) treatment and purification with an RNEasy kit (Qiagen). Embryo trunks were sectioned at 16µm on a cryostat (Leica CM1900) and stored at -20ºC. Prior to in situ hybridisation, slides were defrosted for 1hr at RT. The RNA in situ probe was denatured at 72ºC for 2 minutes and kept on ice. 100µl of hybridisation buffer containing 1ng/µL diluted riboprobe/slide was added to slides and kept in humidified chamber containing formamide overnight at 65ºC. Slides were washed 3 × 30 mins at 65ºC in Wash Buffer (50% Formamide, 5% 20x SSC), then 3x 30mins washes in MABT (Maleic Acid Buffer + 0.1% Tween-20) at RT. Slides were blocked with 300µL Blocking Solution (Blocking Reagent, Sheep Serum, MABT) and kept in humidified chamber at RT for 2 hrs. 75µL of anti-DIG antibody diluted in Blocking Solution was added to slides followed by overnight incubation at RT in a humidified chamber. Slides were washed 4x 20min in MABT followed by 2x 10mins in Alkaline Phosphatase Staining Buffer (4M NaCl, 1M MgCl2, 1M Tris pH 9.5). Slides were then stained with 95µL staining solution (NBT, BCIP, Alkaline Phosphatase Staining Buffer), coverslipped, and kept in the dark at RT overnight. Staining solution was removed by washing 3x 5mins in PBS. Slides were fixed with 300µL 4% PFA for 1hr in a sealed container. Fixative was washed off with 3x 10min PBS washes, and 50µL Mowiol added to each slide for mounting with coverslip. Slides were analysed using brightfield microscopy on Nikon Eclipse Ti Microscope using ND2 Elements software (Nikon).

### qRT-PCR

RNA was extracted from flash frozen embryo heads by using Trizol. Briefly, heads were homogenised in 500uL Trizol. 100uL chloroform was then added,and centrifuged at 6000xg for 30mins. The aqueous layer was removed and an equal amount of 70% EtOH added. The solution was then loaded onto an RNEasy spin column and centrifuged at 13000 rpm for 1 minute. The column was washed with 2x Buffer RLT (Qiagen) and purified RNA eluted in 30µL of RNAse free H20, and stored at -20°C. RNA samples were converted to cDNA using AB Systems High Capacity RNA to cDNA Kit. SYBR Fast standard protocols were used for qPCR with samples run in quadruplicate. B-actin (F-CTGCCTGACGGCCAGG, R- GATTCCATACCCAAGAAGGAAGG) was used to normalise cDNA levels across samples, and *Nes* primers used to measure expression levels across timepoints and samples (F-GCTTCTCTTGGCTTTCCTGA; R- AGAGAAGGATGTTGGGCTGA). Prism software was used for the statistical analysis of qPCR data. Unpaired t-tests were performed to determine if WT *Nes* expression was significantly different from enhancer deleted lines at each timepoint.

## Results

### Generating a *Nes* Enhancer Deletion Mouse Model

To investigate the role of the *Nes* enhancer in directing endogenous expression *in vivo*, we generated an enhancer deletion mouse model using CRISPR/Cas9 mutagenesis. Two gRNAs flanking the *Nes* enhancer were microinjected into mouse zygotes with Cas9 mRNA. 21 founder mice were generated with a range of deletions that partially or completely deleted the *Nes* enhancer. We selected a single founder animal containing both 255bp and 208bp deletion alleles that encompassed all SOXB1 binding sites identified in the ChIP-Seq analysis (Figure 1A). Independent lines were generated for each deletion (referred to hereafter as - 255 and -208). Heterozygous and homozygous -255 pups and embryos were generated at expected ratios, from -255/WT breeding pairs indicating that viability was not compromised by the deletion mutation (Figure 1B,C). No morphological abnormalities were identified in either line indicating that the enhancer deletion did not overtly impact development.

**Figure 1:**
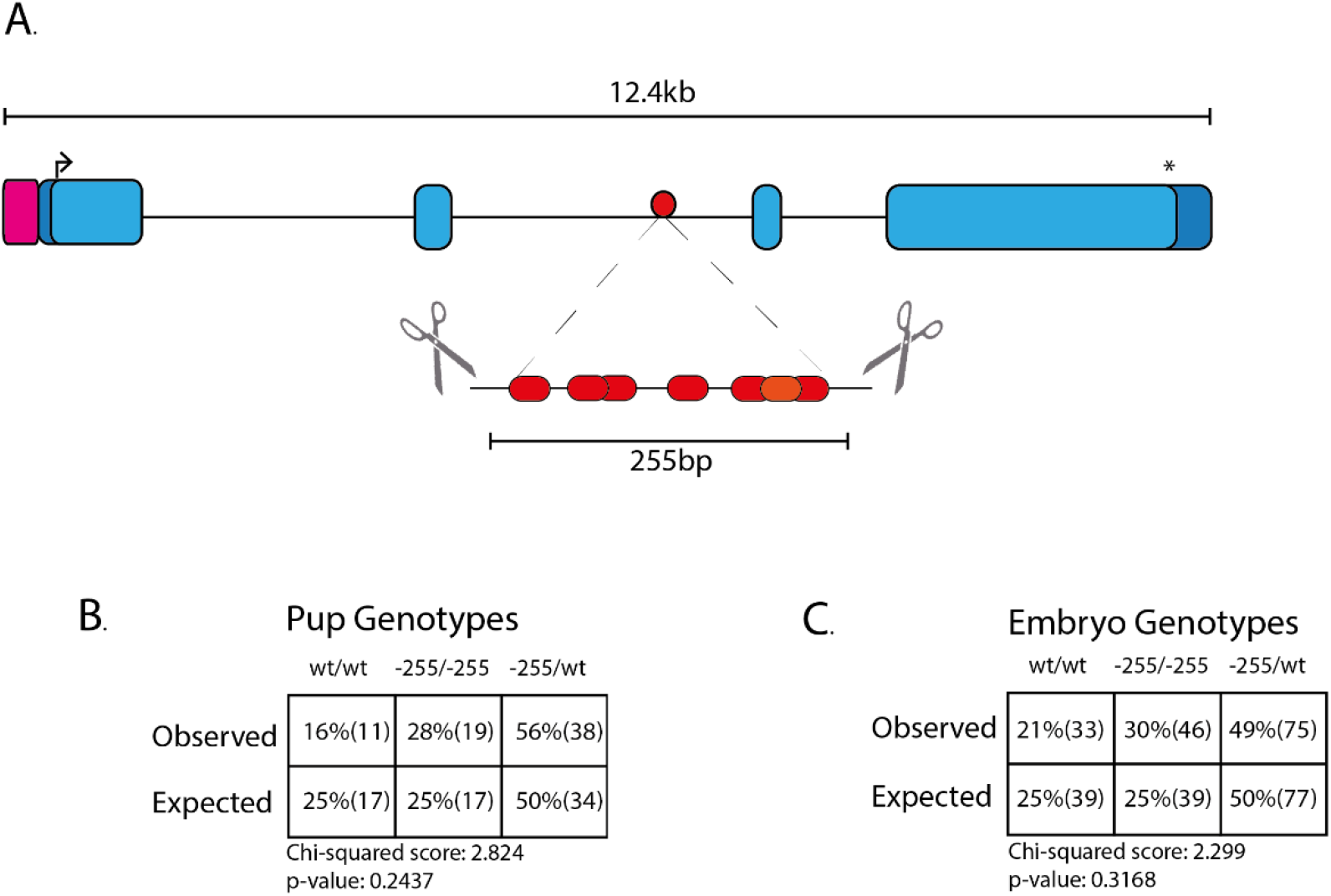
Generation of *Nes* Enhancer deletion (−255/-255) mouse line. A. Guide RNAs (scissors) were designed to flank the six SOXB1 sites (red) and the POU site (yellow) within the second intron of the Nestin gene. Arrow indicates start codon, asterisk indicates stop codon, pink square denotes promoter region and dark blue, the 5’ and 3’ UTRs. The 255bp deletion encompasses all SOXB1 sites identified by ChIP-seq. Observed/Expected tables of both live pups born (B) and transient embryos (C) indicate that there is no embryonic lethality in heterozygous or homozygous animals.

### *Nes* mRNA expression is reduced in enhancer deletion mice

To determine the impact of enhancer deletion on *Nes* expression, qPCR was performed on - 255 homozygous whole embryos (8.5 dpc) and embryonic heads (9.5 dpc-15.5 dpc; Figure 2A). No significant difference in *Nes* expression was detected in mutant embryos at 8.5 dpc. However, from 9.5 dpc significantly reduced levels of *Nes* mRNA were detected in the embryonic cranium. Notably, the greatest reduction in *Nes* expression was detected at 10.5 dpc, with mutant embryos expressing just 18% of *Nes* mRNA compared with WT controls. From 11.5 dpc, a gradual increase in expression was detected in mutants which by 15.5 dpc had recovered to 60% of wild type expression. A reduction in *Nes* expression was also observed in -208 homozygotes at 11.5 dpc (Supplementary Fig. 1).

**Figure 2:**
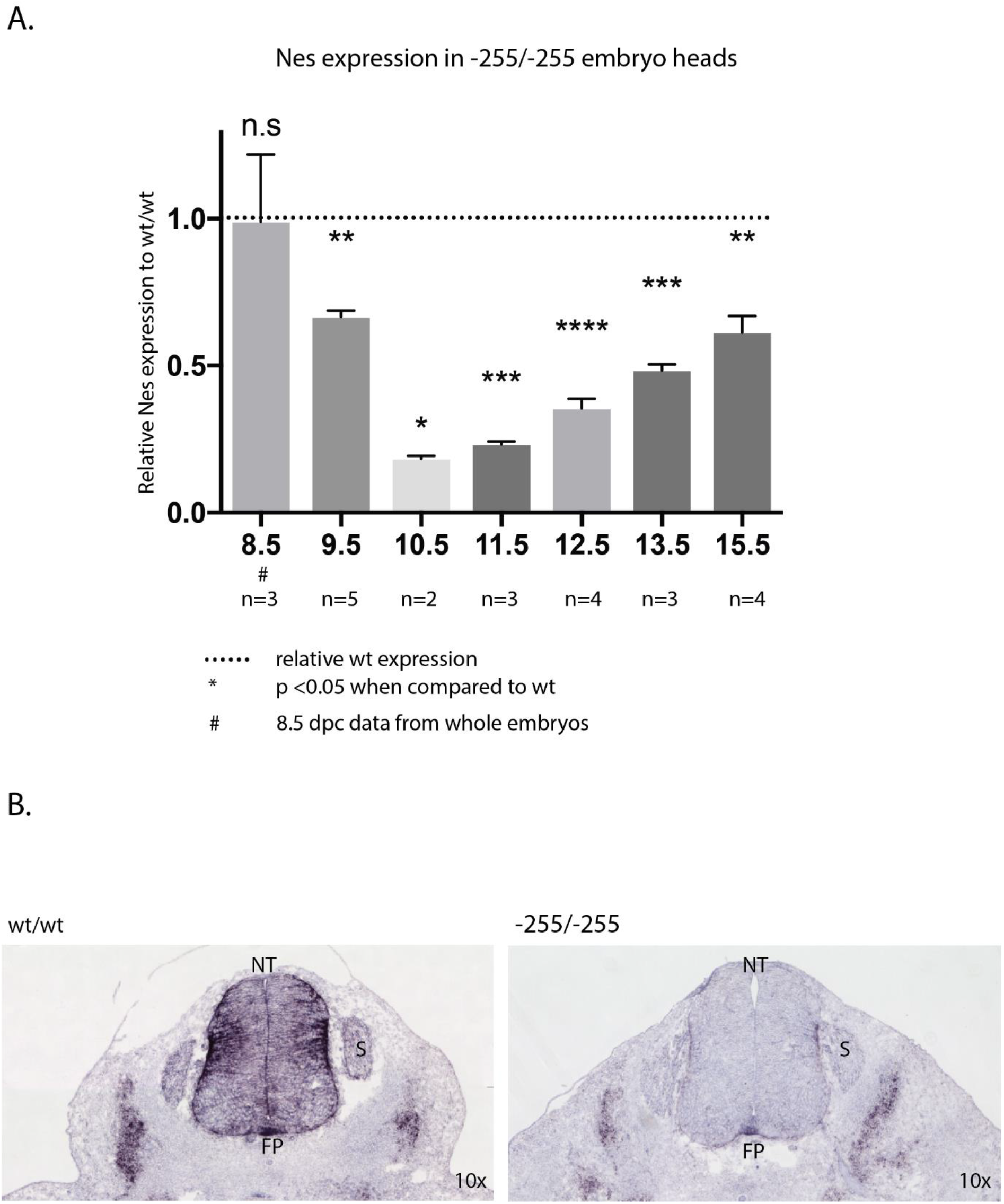
Reduction of Nestin mRNA during embryonic development. A. Analysis of embryonic heads from aged 8.5 to 15.5 by qRT-PCR. All values are normalized to WT samples of the same developmental stage. Due to size constraints, whole 8.5 dpc embryos were used rather than embryonic heads. *Nes* expression is significantly reduced in - 255/-255 embryos from 9.5 dpc-11.5 dpc. * indicates p-value <0.05, ** indicates p-value <0.01, *** indicates p-value <0.001, **** indicates p-value <0.0001. Error bars represent the standard deviation of the mean. B. *In situ* hybridization of *Nes* mRNA at 11.5dpc. Robust *Nes* expression is detected throughout the wild type neural tube.

Next, we determined the spatial impact of enhancer deletion on *Nes* expression in the developing CNS (Figure 2B). For this experiment we analysed the spinal cord at 11.5 dpc as *Nes* is robustly expressed in a stereotypical pattern throughout the trunk at this stage due to the abundance of neural progenitors (Dahlstrand et al. 1995). In situ hybridization was performed on the trunk sections of wildtype (WT) and homozygous enhancer deletion (−255) embryos. As expected, expression of *Nes* was detected throughout the spinal cord, with the highest levels confined to the lateral regions and the floor plate. In contrast, the spinal cord of enhancer-deleted embryos was virtually devoid of *Nes* mRNA, except for restricted expression in the floor plate and the lateral regions. Notably, lateral expression in the mesoderm was not noticeably diminished in mutant embryos, consistent with the neural-specific activity of the *Nes* enhancer in transgenic mice. Thus, deletion of the *Nes* neural enhancer results in a striking reduction in the level and extent of *Nes* expression.

### NES protein expression is reduced in enhancer deleted mice

As *Nes* mRNA expression was significantly reduced in both the embryonic head and neural tube, we performed protein expression analysis in these regions to determine whether NES was similarly reduced. Both head and trunk transverse sections were prepared from WT and homozygous -255 embryos and co-stained with anti-NES (trunk and brain) and anti-SOX3 antibodies (trunk). We hypothesised that as the enhancer is controlled by the SOXB1 proteins binding to the region, that we would see little to no NES expression throughout the SOX3 expressing zones of the neural tube and brain.

The WT brain sections show the telencephalon is densely stained for Nestin, showing a long filamentous structure without nuclear staining (Figure 3A). In the homozygous enhancer deletion however, there are obvious differences in the staining pattern of the NES protein, as it is duller throughout the telencephalon, and shows regions of high reactivity that appear to be within the vasculature.

**Figure 3:**
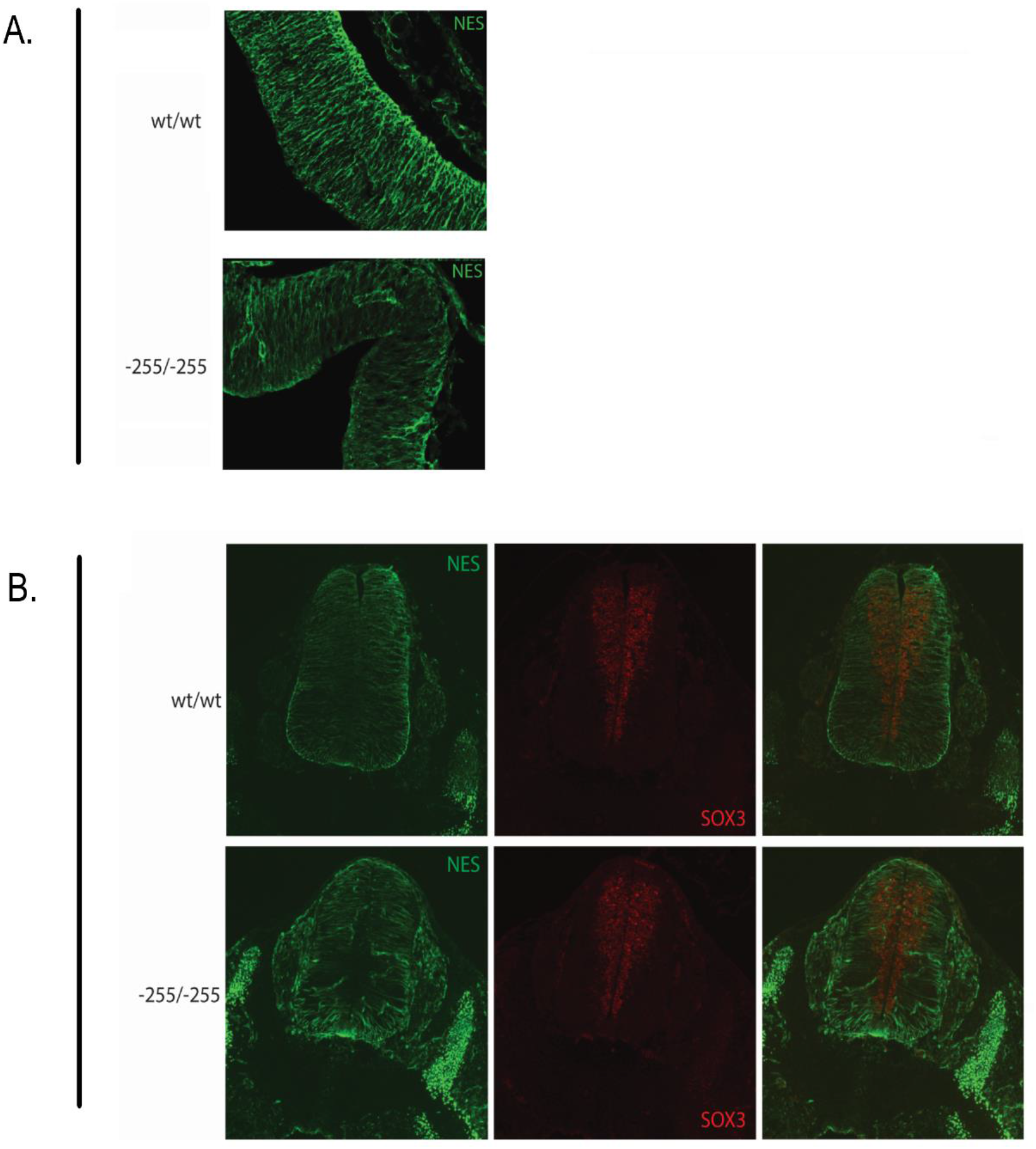
Immunohistochemical analysis of brain and trunk sections. Wildtype(WT/WT), heterozygous (−255/WT) and homozygous (−255/-255) transverse cortex (A) and trunk (B) sections labelled with anti-SOX3 and anti-NESTIN antibodies. The NES signal is decreased in -255/-255 sections, while the SOX3 remains consistent across genotypes.

This experiment was repeated using neural tube sections, with SOX3 and NES antibodies, and similar results were obtained (Figure 3B). The WT embryos exhibit smooth filamentous NES staining from the lateral edges towards the midline. In contrast, NES signal in the -255/-255 embryos was weaker, particularly within the periluminal SOX3-positive region. Taken together with the mRNA expression analysis, these data confirm that NES expression is reduced in the developing nervous system of enhancer-deleted embryos.

### Ectopic Nestin expression in vasculature of enhancer deleted embryos

Whilst analysing -255/-255 embryos for NES protein expression, we noted specific staining in discrete structures within the neural tube and cortex that appeared to be the developing vasculature. Notably, this signal was not present in WT or heterozygous embryos. To further investigate this finding, we co-stained 10.5 dpc embryo heads with antibodies to the endothelial cell marker CD31 and NES (Figure 4A). Images captured using an inverted fluorescence microscope indicated colocalisation of NES and CD31 in -255/-255 embryos but not in WT controls. Additional analysis using confocal microscopy revealed widespread expression of NES in endothelial cells lining the developing vasculature of -255/-255 embryos. In contrast, NES expression was rarely detected in WT endothelial cells. Thus, deletion of the *Nes* neural enhancer induces ectopic expression in endothelial cells.

**Figure 4:**
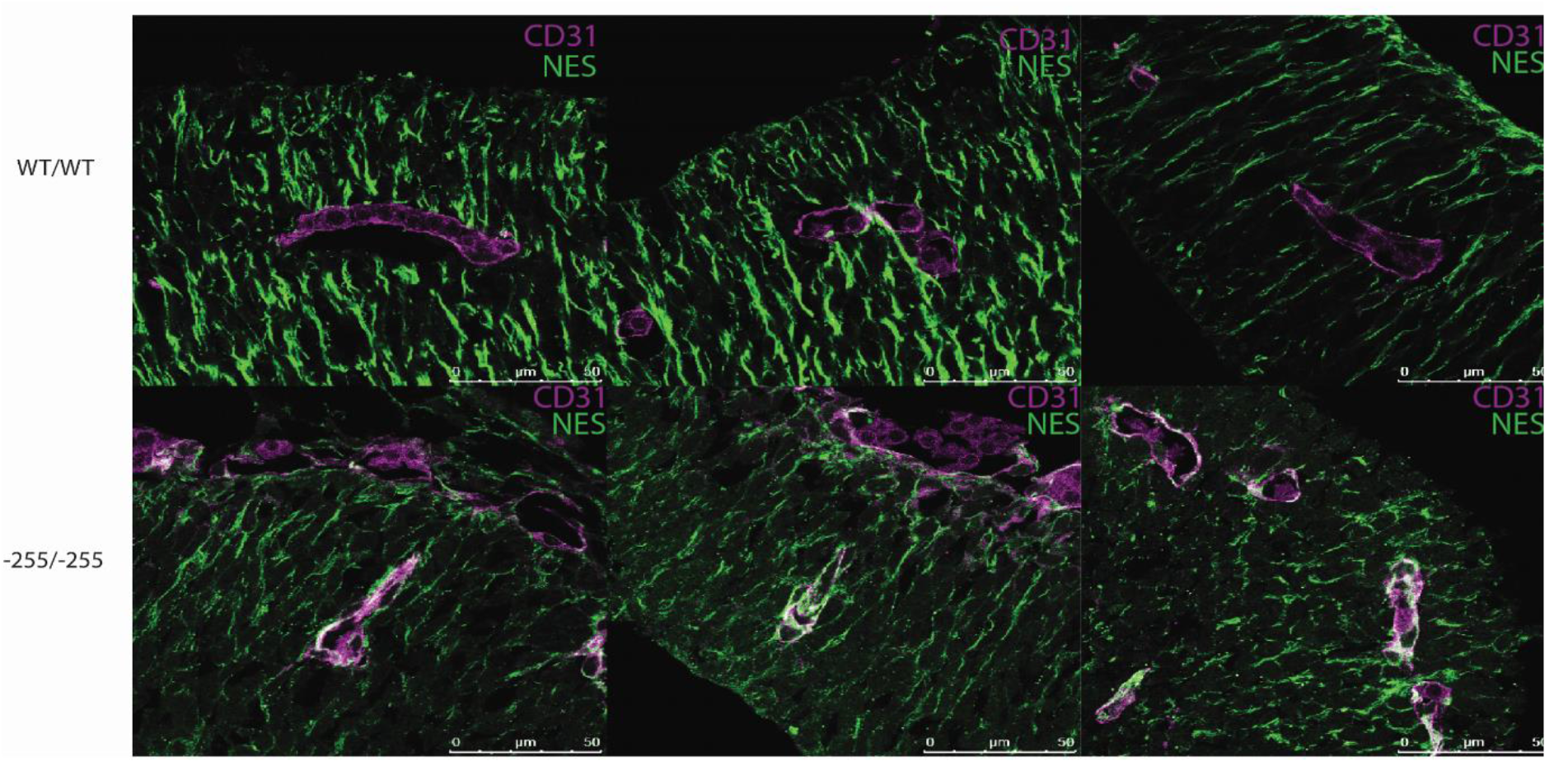
Immunohistochemical analysis of Nestin reactivity within vasculature of 10.5 dpc cortex sections. Confocal microscopy of WT/WT and -255/-255 10.5 dpc cortex sections labelled with NESTIN and CD31 to mark epithelial cells of the vasculature. Co-localisation (white) of Nestin and CD31 is apparent in the -255/-255 sections, and not seen in WT sections.

### *Nes* is not required for CNS development

It has previously been reported that deletion of *Nes* causes extensive cell death in the developing CNS and embryonic lethality at approximately 8.5 dpc (7). Given that -255/-255 mutants do not exhibit overt developmental defects, it appears that the level of *Nes* in these enhancer-deleted embryos exceeds the threshold required for normal development. We were therefore interested in assessing whether further reduction of *Nes* levels in -255/KO compound heterozygous embryos would compromise CNS development. To generate *Nes* knockout mice, we employed a dual gRNA deletion strategy (Figure 5). The proximal gRNA targeted exon 1 immediately downstream of the start codon and the distal gRNA cut immediately upstream of the stop codon in exon 4. The rationale for this approach was that null alleles could be generated via frameshifting indels at the proximal cut site or from deletion of the ∼8.7 kb intervening sequence between the proximal and distal cut sites. This approach also provided the necessary alleles for the *trans* enhancer interaction experiment (see below). PCR genotyping indicated that four of the six founder animals contained at least one large deletion allele. Sanger sequencing confirmed that the founder used for subsequent breeding carried the expected 8672 bp deletion that encompassed almost all of the coding region and introns of the *Nes* gene, including the neural enhancer in intron 2.

**Figure 5:**
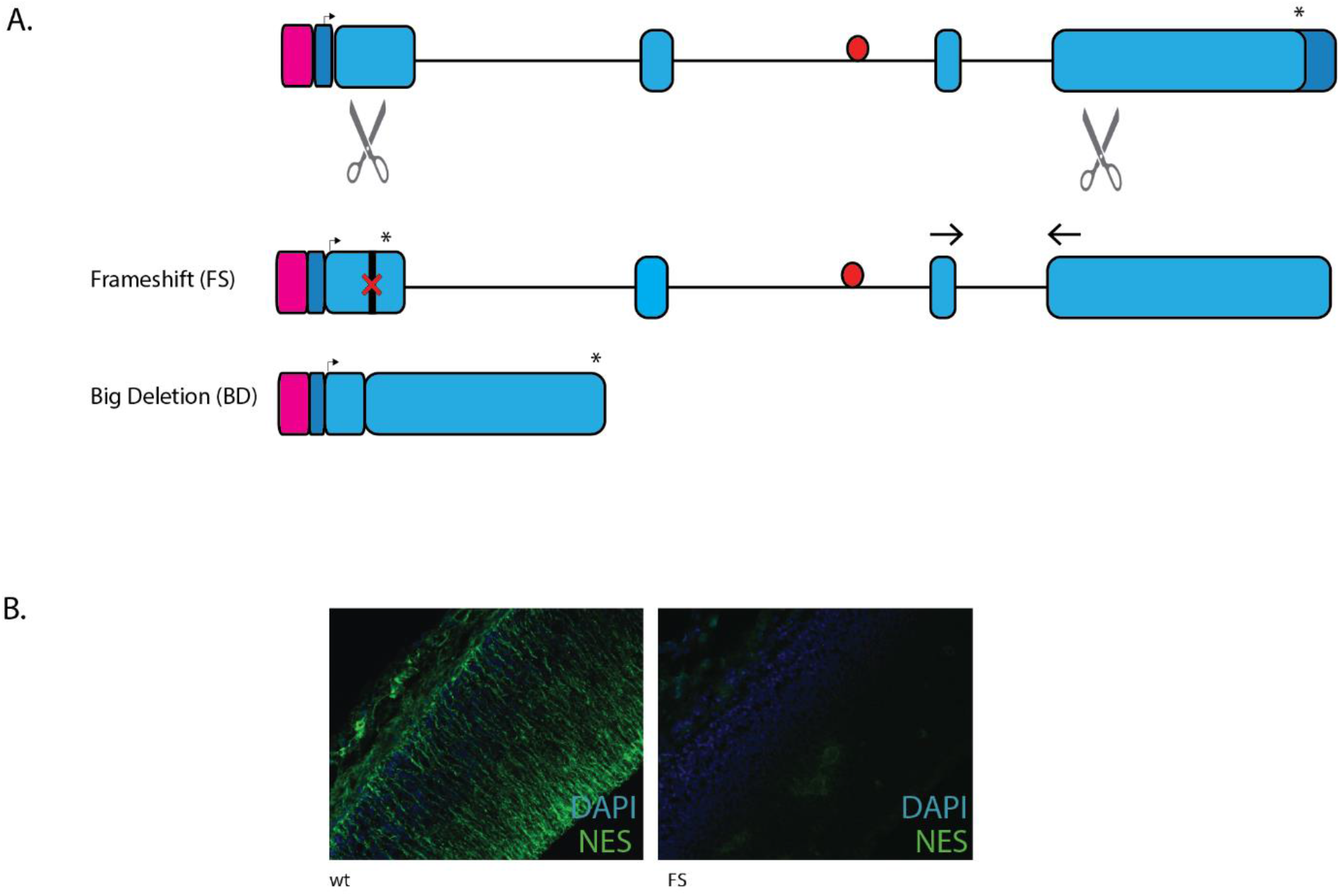
Generation of *Nes* null mouse lines. A. CRISPR guide sequences (scissors) designed to cut within exon 1 and exon 4 of the *Nes* gene. The FS allele generated a frameshift mutation at codon 50, while the BD allele removed the 8.7 kb of intervening sequence. qRT-PCR primers indicated by the arrows amplify the FS, WTand -255 alleles. Pink box indicates the promoter, dark blue is 5’ and 3’ UTR, pale blue is coding regions, red circle is *Nes* enhancer, arrow is transcriptional start site and asterisk is the stop codon. B. Immunohistochemical staining with NES antibody on ‘frameshift’ embryonic cortex shows no detectable protein product.

**Figure 6:**
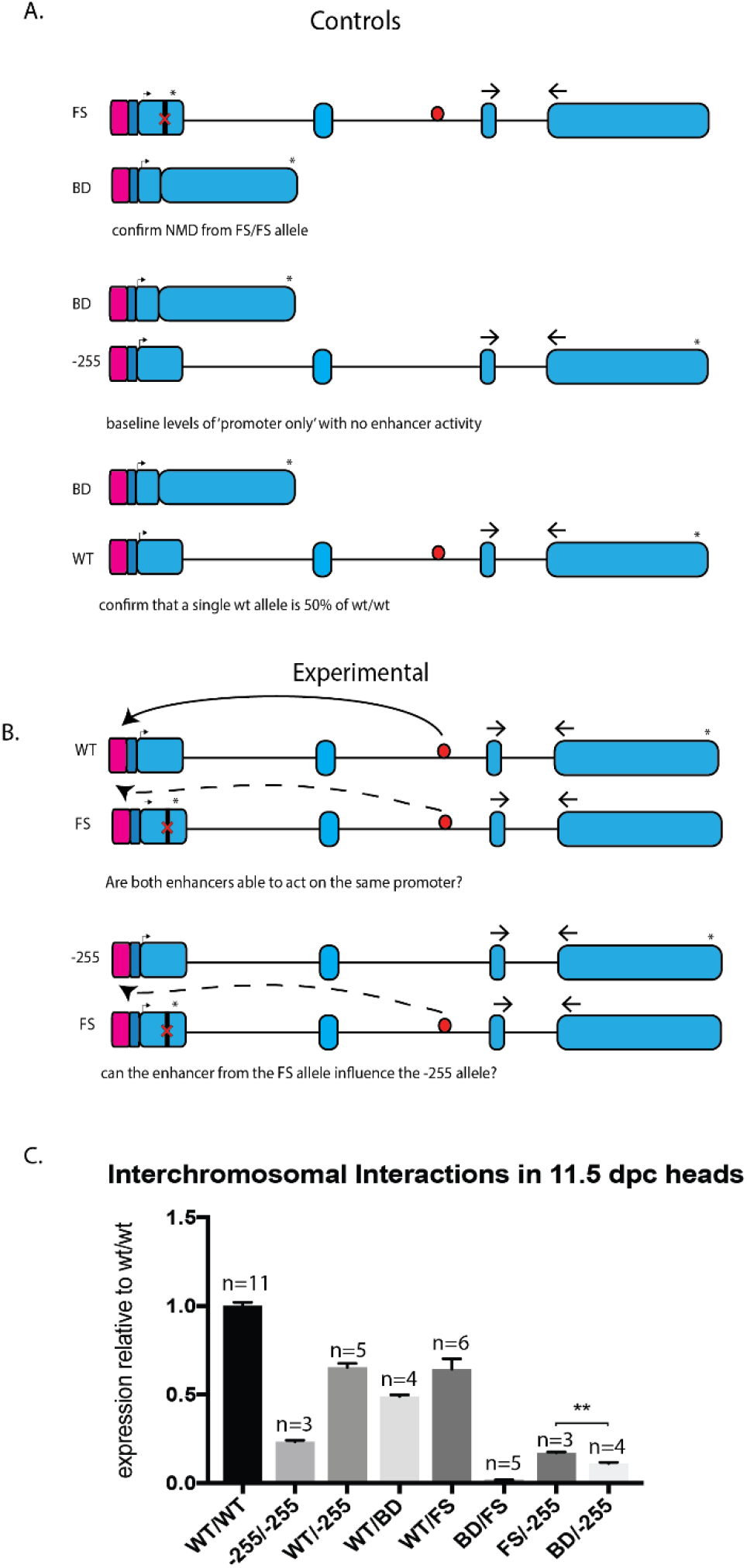
Interchromosomal Interactions of the Nestin enhancer and promoter. A. Control crosses to determine trans interactions. The first will determine if nonsense mediated decay occurs in the ‘big deletion’ (BD) allele. The second determines the baseline level of ‘promoter only’ activity when no enhancer is present. The third will confirm ifa single allele produces exactly half of the total WT mRNA. qRT-PCR primers indicated by the arrows amplify the FS, WT and -255 alleles. Pink box indicates promoter, dark blue is 5’ and 3’ UTR, pale blue is coding regions, red circle is *Nes* enhancer, arrow is transcriptional start site and asterisk is the stop codon. B. Experimental workflow to determine *trans* interactions. The first will determine whether both copies of the *Nes* enhancer are capable of influencing only one functional promoter. The second will determine if an enhancer on one allele can compensate for the loss of the enhancer on another allele. qRT-PCR primers indicated by the arrows amplify the FS, WT and -255 alleles. Pink box indicates promoter, dark blue is 5’ and 3’ UTR, pale blue is coding regions, red circle is *Nes* enhancer, arrow is transcriptional start site and asterisk is the stop codon. C. The qPCR results of the above experimental crosses and embryo analyses on 11.5 dpc heads. By using various mating of FS, BD, WT and -255 alleles embryonic heads were analysed for changes in Nestin gene expression. The BD/-255 produces significantly less Nestin mRNA that the FS/-255, indicating that the presence of a single enhancer on one chromosome can interact with the promoter of another. Unpaired t-test between FS/-255 and BD/-255 show p-value of 0.0011, other t-test results not shown for clarity. Error bars represent the standard deviation of the mean.

This null allele, *Nes* g.54_4518/p.L19_V1506del/p.L19fsX, termed BD, encodes only the first 18AA of the NES open reading frame, and a frameshift causes the last 30AA of exon 4 to be incorrect. This founder also carried an 8bp frameshifting deletion, g.50_57del/p.R17fsX75, termed FS, at the proximal cut site that terminated the protein after 13 amino acids. Breeding colonies for each mutation were generated. Surprisingly, BD and FS homozygous mice were viable and did not exhibit any obvious phenotype or developmental defects. Compound heterozygous FS/BD mice were also phenotypically normal. To confirm that NES protein was not generated from the FS allele, we stained FS/FS embryonic brain sections with anti-NESTIN antibody. In contrast to WT control tissue, no expression was detected in mutant tissue. We therefore conclude that *Nes* is not required for CNS development or viability.

### *Trans* Interactions of the *Nes* neural Enhancer

While enhancers are generally considered to be *cis*-regulatory elements, previous studies have provided evidence for interchromosomal *trans* interaction between enhancers and their cognate promoters (18). To investigate possible interchromosomal activity of the *Nes* neural enhancer *in vivo*, we used qPCR to measure allele-specific expression in a series of compound heterozygous embryos. For this experiment, we exploited the presence of the *Nes* enhancer in the FS allele but not the BD allele. Thus, any difference in *Nes* expression from the WT allele in FS/WT and BD/WT embryos would reflect *trans* activity of the (FS) *Nes* enhancer. Similarly, any difference in *Nes* expression from the -255 allele in FS/-255 and BD/-255 embryos would reflect *trans* activity of the (FS) Nestin enhancer. No detectable signal was generated from FS/BD embryos indicating that *Nes* mRNA is not generated from either null allele (presumably due to nonsense mediated decay for the FS allele). -255/-255 *Nes* expression was 23% of WT expression, consistent with our previous analysis (Fig. 2A). Comparison of FS/-255 and BD/-255 expression revealed significantly higher expression in the former (17% vs 11%; p< 0.01). Similarly, WT/FS *Nes* expression was higher than WT/BD, although this did not reach significance (64% vs 49%; p< 0.07). Together, these data suggest that the *Nes* enhancer may function in *trans*.

## Discussion

While enhancers are routinely used to drive spatio-temporally restricted expression of heterologous genes, their functional role in coordinating cognate gene expression remains poorly understood. Using CRISPR/Cas9 technology, we show that deletion of the *Nes* neural enhancer has a profound impact on endogenous *Nes* expression in the developing nervous system. Our data indicate that this region also contains a repressor element that inhibits expression in endothelial cells, underlining the ability of deletion analysis to identify both positive negative regulatory interactions.

*Nes* is expressed within the incipient neural progenitor cells during early embryogenesis and is maintained during expansion of this cell population. Upon differentiation, *Nes* is downregulated and is replaced by other members of the intermediate filament family (19). At 8.5 dpc, *Nes* expression is not significantly different in -255 homozygous embryos indicating that the neural enhancer is not functionally required for initiation of *Nes* expression. Given that there is robust SOXB1 expression in neuroprogenitors at this stage, it appears that putative binding of these factors to the *Nes* enhancer at this stage is not required for expression but may nevertheless play a role in maintaining the locus in an “open for business” conformation (20). From 9.5 dpc, *Nes* expression is significantly lower in -255 homozygous embryos. Indeed, at 9.5 dpc, the neural enhancer is required for approximately 80% of the *Nes* expression within the head. The activity of the enhancer remains functionally significant until at least 15.5dpc, although the differential between - 255 and WT expression becomes less pronounced, suggesting that other neural enhancer(s) have increasingly important roles as the nervous system develops. It is interesting to compare our data with other recently published examples of developmental enhancer deletion. A study of single conserved limb enhancers showed that 90% of these had no impact on cognate gene expression (21), as well as highly conserved ß-globin enhancers (22). Computational analyses have predicted that most (96%) of enhancers within the genome are tolerant to LoF mutations, and that many essential genes will have a degree of enhancer redundancy (23). In the vast majority of examples, deletion of a single conserved enhancer element has no impact on cognate gene expression (21, 22). The Nestin enhancer therefore appears to be unusual in having such a profound impact on *Nes* expression during early embryonic development. The mechanism that underpins this unusually high activity remains to be determined, as well as elucidating other *Nes* regulatory elements that are responsible postnatally.

The expression of *Nes* mRNA throughout the neural tube is considerably affected in mutants lacking the 255bp enhancer, as seen in both in situ hybridization and qRT-PCR experiments. Decreased protein reactivity is seen in the -255 embryos, however the staining is still present throughout the neural tube where the mRNA is not visualized. This is possibly due to very low levels of Nestin expression within these cells, undetectable through in situ hybridization. It is also expected that *Nes* expression is controlled by other transcription factors other than SOX or POU proteins that are expressed within non-SOX/POU regions.

An unexpected finding of this study was that deletion of the *Nes* enhancer resulted in ectopic expression within the vasculature. Previously, NES has been reported to be expressed within the vasculature of different tissues such as developing kidneys, and also shown to be upregulated within vasculature following focal cerebral ischemia (24-26), indicating a role in development and repair. Whilst the mechanism is unclear, it appears that the -255 deleted region also contains a repressor element that prevents NES expression in developing vasculature. While further studies are required to determine the protein-sequence interaction(s) that mediate this repressor activity, it is worth noting that unmasking of repressor elements cannot be achieved using traditional enhancer activity assays such as transgenic reporter analysis, highlighting the utility of the enhancer deletion approach we have employed.

Within the literature there are conflicting reports as to whether *Nes* is an essential gene in mice. The first reported *Nes* null line generated showed early embryonic lethality, and despite two further *Nes* null lines showing homozygous viability, *Nes* is often cited as an essential gene (6, 7, 27, 28). Through generation of two independent CRISPR KO mouse lines, we have shown that NES null mice are viable and do not exhibit overt deleterious phenotypes, consistent with two previous reports (6, 27). In contrast, the NES null mice reported by Park, Xiang (7) exhibit embryonic lethality at 8.5dpc due to apoptosis of neural tube cells. The reason for this inconsistency remains unclear. Although not explored in this study, mild phenotypes such as impaired motor coordination (6) in KO mice suggest that NES function cannot be entirely replaced by other members of the intermediate filament family.

It is often assumed that all enhancers only act in *cis* to regulate cognate gene expression. However, it remains unclear whether some enhancers can also function in *trans* to activate cognate target gene(s). *Trans* enhancer interactions or transvection is well characterised in *Drosophila* (2, 29) but has rarely been observed in mammals (3). Utilising a genetic approach, we provide evidence that the Nestin enhancer can undergo functional *trans* interactions *in vivo*. While the effect is relatively weak, these data raise the possibility that transvection of developmental enhancers may be more common than is currently recognised. Further investigation using chromatin capture technology would be beneficial in characterising these putative *trans* interactions. While this technology has been successfully used to characterise *trans* interactions within *Drosophila*, this has not yet shown interactions within the mouse (30, 31).

