## supplemental figure for "The Nestin neural enhancer is essential for normal levels of endogenous Nestin in neuroprogenitors but is not required for embryo development"

A.

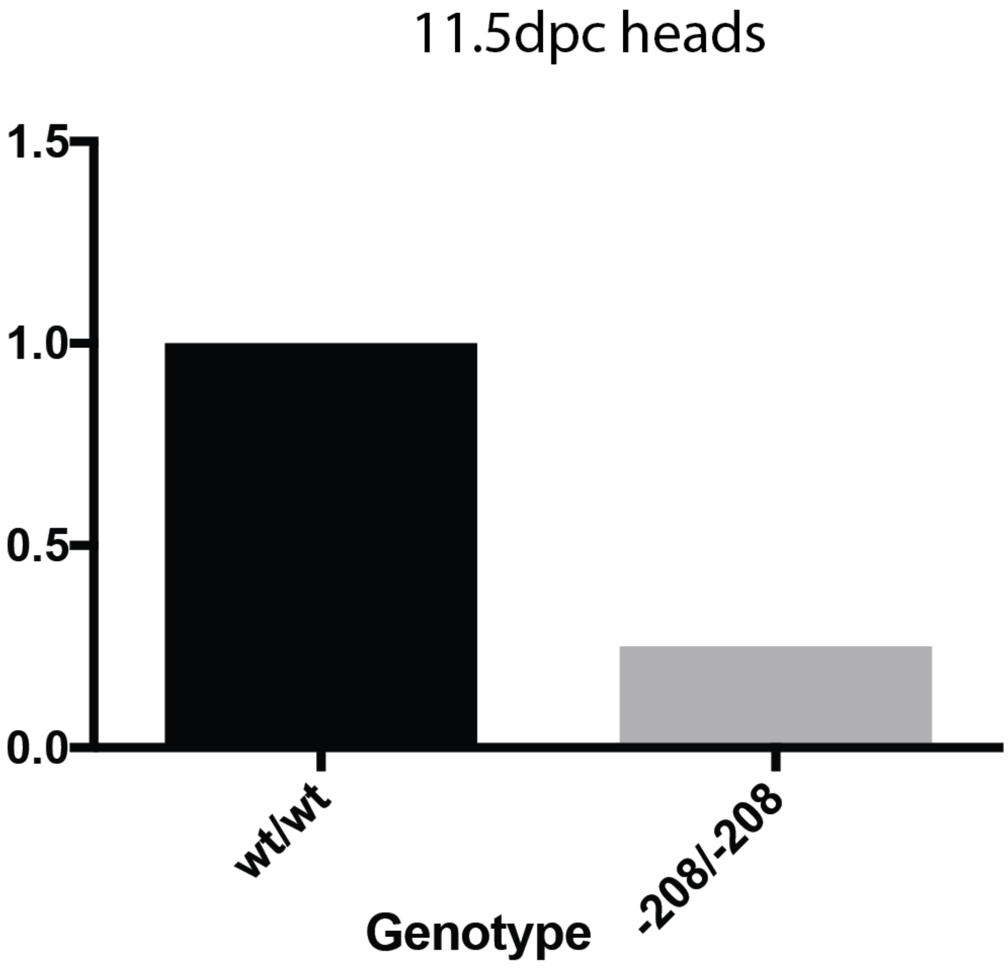


**Supplementary Figure 1**

The -208 Nestin enhancer deletion line shows a reduction in *Nes* expression in 11.5 dpc embryonic heads similar to that of the -255 line (n=2 embryos).
